# Identification of cell barcodes from long-read single-cell RNA-seq with BLAZE

**DOI:** 10.1101/2022.08.16.504056

**Authors:** Yupei You, Yair D.J. Prawer, Ricardo De Paoli-Iseppi, Cameron P.J. Hunt, Clare L. Parish, Heejung Shim, Michael B. Clark

**Affiliations:** School of Mathematics and Statistics/Melbourne Integrative Genomics, The University of Melbourne, Parkville, VIC, 3010, Australia; Centre for Stem Cell Systems, Department of Anatomy and Physiology, The University of Melbourne, Parkville, VIC, 3010, Australia; The Florey Institute of Neuroscience and Mental Health, The University of Melbourne, Parkville, VIC, 3010, Australia

## Abstract

Single-cell RNA sequencing (scRNA-seq) has revolutionised our ability to profile gene expression. However, short-read (SR) scRNAseq methodologies such as 10x are restricted to sequencing the 3’ or 5’ ends of transcripts, providing accurate gene expression but little information on the RNA isoforms expressed in each cell. Newly developed long-read (LR) scRNA-seq enables the quantification of RNA isoforms in individual cells but LR scRNA-seq using the Oxford Nanopore platform has largely relied upon matched short-read data to identify cell barcodes and allow single cell analysis. Here we introduce BLAZE (Barcode identification from long-reads for AnalyZing single-cell gene Expression), which accurately and efficiently identifies 10x cell barcodes using only nanopore LR scRNA-seq data. We compared BLAZE to existing tools, including cell barcodes identified from matched SR scRNA-seq, on differentiating stem cells and 5 cancer cell lines. BLAZE outperforms existing tools and provides a more accurate representation of the cells present in LR scRNA-seq than using matched short-reads. BLAZE provides accurate cell barcodes over a wide range of experimental read depths and sequencing accuracies, while other methodologies commonly identify false-positive barcodes and cell clusters, disrupting biological interpretation of LR scRNA-seq results. In conclusion, BLAZE eliminates the requirement for matched SR scRNA-seq to interpret LR scRNA-seq, simplifying procedures and decreasing costs while also improving LR scRNA-seq results. BLAZE is compatible with downstream tools accepting a cell barcode whitelist file and is available at https://github.com/shimlab/BLAZE.

## Background

Single-cell transcriptomics has become a widely accessible and popular means of profiling gene expression at single-cell resolution. The applications of single-cell RNA sequencing (scRNA-seq) are broad, ranging from identification of cell and tissue types, tracking developmental trajectories, and assessing system heterogeneity [1]. However, short-read (SR) scRNA-seq methodologies lack the ability to accurately identify RNA isoforms. Droplet based platforms such as the popular 10x platform [2] are restricted to sequencing the 3’ or 5’ ends of transcripts, providing accurate gene counts but little information on RNA splicing or the RNA isoforms expressed in each cell [3]. Alternative methods, such as Smart-seq3, sequence all parts of transcripts but are still constrained by short sequencing read lengths, which largely prevents the accurate reconstruction of RNA isoforms longer than 1 kb [4].

The recent development of long-read (LR) single-cell sequencing methods has laid the foundation for a more in-depth analysis of isoforms for single cells [5]. LR scRNA-seq methods have been developed using both the PacBio and Oxford Nanopore Technologies (ONT) platforms, allowing for the discovery and quantification of full-length RNA isoforms in single cells [6-19].

Two key steps in enabling scRNA-seq analysis are the identification of cell barcodes, which denote which cell a read is from, and unique molecular identifiers (UMIs), which allow removal of PCR duplicates and more accurate counting of gene and isoform expression. A limitation of most LR scRNA-seq methodologies is that they require matched SR scRNA-seq for the identification of cell barcodes and/or UMIs, particularly those using nanopore sequencing due to its higher error rate [8, 9, 12, 14-17, 19]. The addition of matched SR data adds technical complications for library construction and significantly increases both the time and cost needed to produce these data sets. Furthermore, the requirement for matched SR data can also greatly decrease the proportion of usable long-reads [9]. Other methods for nanopore LR scRNA-seq have been reported that do not require the addition of matched short-reads. However, these methods are either very low throughput [10], require bespoke reagents and are incompatible with existing 10x workflows [13], or trade higher accuracy for lower read depth [18]. Therefore, a method which requires only nanopore LRs and is compatible with existing workflows is required [15]. Recently, ONT released the Sockeye pipeline (https://github.com/nanoporetech/Sockeye) to perform LR-only scRNA-seq analysis, including barcode and UMI identification. However, the performance of Sockeye is yet to be determined.

Here we introduce BLAZE (**B**arcode identification from **L**ong-reads for **A**naly**Z**ing single-cell gene **E**xpression), which accurately identifies 10x cell barcodes using only nanopore LR scRNA-seq data. In combination with the existing FLAMES pipeline [15], BLAZE eliminates the requirement for matched SR scRNA-seq, simplifying LR scRNA-seq workflows, reducing sequencing costs and producing improved results. We show that BLAZE performs well across different sample types, sequencing depths and sequencing accuracies and outperforms other barcode identification tools such as Sockeye. We designed BLAZE to seamlessly integrate with the existing FLT-seq - FLAMES pipeline to enable identification and quantification of RNA isoforms and their expression profiles across individual cells and cell-types. Taken together, BLAZE provides a cheaper, simpler and more accurate means to profile transcript-level changes in LR scRNA-seq data sets.

## Results

### Single-cell barcode identification with BLAZE

We designed BLAZE for the accurate identification of cell barcodes from Oxford Nanopore long-read libraries generated using the 10x single-cell 3’ gene expression profiling procedure. To enable cell barcode identification from nanopore reads despite their higher error rate, BLAZE performs a three-step procedure (**Fig. 1A**, see methods for further details). First, BLAZE identifies the likely position of the cell barcode and extracts the putative barcode sequence. The 16 nt barcode and 10-12 nt UMI are located between the adapter and polyT sequences. BLAZE locates the cell barcode in each read by identifying the probable adaptor and polyT regions. The 16 nt sequence immediately downstream of the adaptor is defined as the “putative barcode”. BLAZE discards putative barcodes that do not appear in the list of all possible 10x barcodes because these cannot represent true barcodes. Next, BLAZE selects high-quality putative barcodes whose sequences are less likely to contain basecalling errors. Specifically, BLAZE filters out barcodes with a minimum quality score (denoted as “minQ”) of less than 15 across the 16 bases that comprise the putative barcode (**Additional File 1: Fig. S1**). Finally, BLAZE counts the occurrence of each unique high-quality barcode, ranks them based on those counts, and selects the top-ranked ones as barcodes likely associated with cells using a quantile-based threshold.

**Fig. 1:**
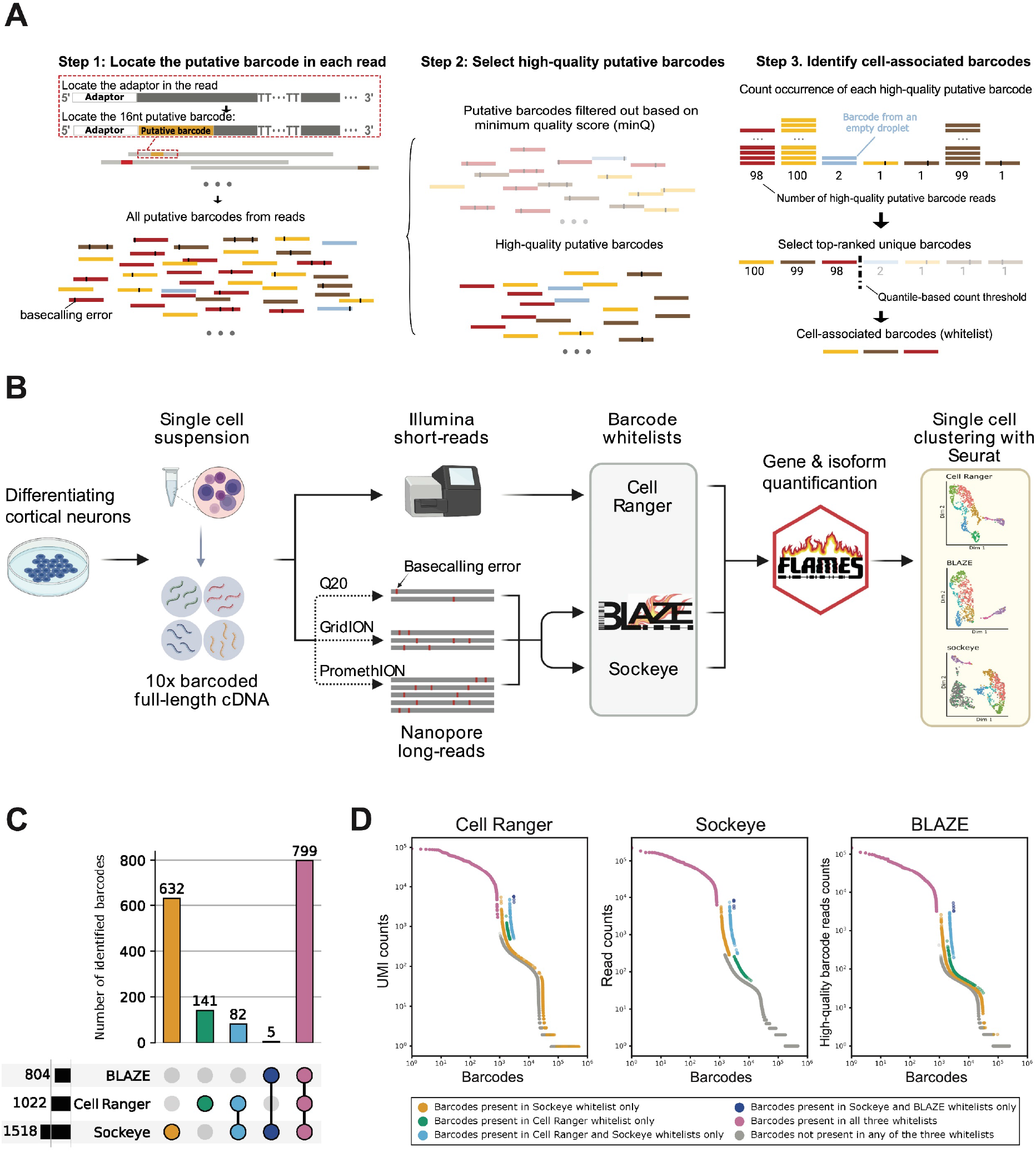
Experimental overview and comparison of identified cell barcodes. **A: BLAZE Workflow.** Step 1: locate putative barcodes by first locating the adaptor in each read. Putative barcodes include those originating from different cells and empty droplets. In schematic putative barcodes with the same colour come from the same original cell/droplet. Black blocks on putative barcodes represent basecalling errors. Step 2: Select high-quality putative barcodes. Bases representing sequencing errors tend to have low quality scores. Putative barcodes with minQ < 15 are filtered out (faded in the figure) and the majority of the remaining putative barcodes are expected to have no errors. Step 3: Identify cell-associated barcodes. BLAZE counts and ranks unique high-quality putative barcodes and outputs a list of cell-associated barcodes whose counts pass a quantile-based threshold. **B: Schematic of experimental design**. Human induced pluripotent stem cells (hiPSC) undergoing cortical neuronal differentiation were dissociated into a single-cell suspension and processed to generate single-cell full-length cDNA. Full-length cDNA was sequenced using both short and long read methods and barcode whitelists generated using Cell Ranger, BLAZE and Sockeye followed by gene and isoform quantification and clustering. Three nanopore sequencing runs were performed on the same cDNA sample, a higher depth PromethION run, a lower depth GridION run and a higher accuracy run using the Q20 protocol on the GridION. **C: Barcode upset plot comparing different whitelists**. The bar chart on the left shows the total number of barcodes found by each tool. The bar chart on the top shows the number of barcodes in the intersection of whitelists from specific combinations of methods. The dots and lines underneath show the combinations. The colours of the combinations are used to distinguish barcodes in figure 1D. **D: Barcode rank plot**. Unique barcodes are ranked based on the counts output by each method and coloured by which method(s) included each barcode in their barcode whitelist(s). The colours for different combinations of methods follow those in figure 1C and barcodes not included in any of the whitelists are in grey. Cell Ranger short-read counts, Sockeye long-read counts and BLAZE long-read counts shown on left, middle and right knee plots respectively. Sockeye and BLAZE analyse the same dataset. Cell Ranger analyses counts from a short-read library, deriving from the same original cDNA. Unique barcodes are ranked on the x-axis based on the number of reads/unique molecules observed for each (y-axis). Shifts on the x-axis are intentionally added to make the dots with different colours non-overlapping. Note that these three methods generate counts in different ways so the three plots have different y-axis labels.

A significant proportion of putative barcodes are expected to be error-free, despite the ∼4-5%, (or ∼2% with higher-accuracy Q20 protocols) median error rate for nanopore reads (**Table 1**). With sufficient per-cell sequencing depth, this means each cell should be supported by large number of high-quality putative barcodes. Therefore, highly-supported barcodes likely represent true cells while poorly-supported barcodes likely represent sequencing errors and/or barcodes associated with empty droplets (**Fig. 1A**). The output of BLAZE is a list of unique cell-associated barcodes (referred to as barcode whitelist), that is input into downstream gene and isoform quantification software in place of a whitelist generated from matched SR sequencing.

**Table 1:** Summary statistics for long-read single-cell data sets.

| Dataset ID | Sequencing platform | Sequencing Kit | Total Reads | Pass reads | Median accuracy of pass reads (%) | Useable* reads with Cell Ranger | Useable reads with BLAZE | Useable reads with Sockeye |
| --- | --- | --- | --- | --- | --- | --- | --- | --- |
| PromethION | PromethION | SQK-LSK110 | 100,941,015 | 61,967,455 | 95.1 | 43,473,150 (70%) | 43,001,188 (69%) | 43,063,370 (69%) |
| GridION | GridION | SQK-LSK110 | 10,371,632 | 7,521,667 | 96.0 | 5,461,630 (73%) | 5,430,433 (72%) | 5,462,816 (73%) |
| Q20 | GridION | SQK-Q20EA | 4,479,225 | 3,423,062 | 97.9 | 2,501,260 (73%) | 2,484,366 (73%) | 2,511,016 (73%) |
| scmixology2 (Tian *et al.* 2021) | PromethION | SQK-LSK109 | 36,927,546 | 25,164,948 | 95.7 | 18,347,232 (73%) | 18,099,553 (72%) | 17,818,239 (71%) |

\*Pass reads that have a valid 10x barcode, a UMI and can be assigned to a barcode in whitelist.

### Experimental workflow to assess the performance of BLAZE

We tested the performance of BLAZE by carrying out matched short and long-read scRNA-seq on ∼1000 human induced pluripotent stem cell (iPSC)-derived neural progenitors undergoing differentiation to the cortical lineage (**Fig. 1B**, see methods). Short-reads were sequenced on an Illumina NOVA-seq to a high median depth of 96,000 reads per cell. SR data were analysed with the Cell Ranger pipeline (10x Genomics) to generate a barcode whitelist that can be directly compared to a whitelist generated from long-reads only. We performed LR scRNA-seq using the FLT-seq protocol [15] and sequenced the sample on a PromethION flowcell. generating ∼62 million pass reads (**Table 1**). In addition to deep PromethION sequencing, we also sequenced the cDNA on the GridION using standard and higher accuracy (Q20) chemistries generating ∼7.5 and ∼3.5 million pass reads respectively (**Table 1**). This enabled us to assess the effects of read depth and variation in read accuracy on the performance of BLAZE and is discussed in greater detail below. We also compared BLAZE to Sockeye (https://github.com/nanoporetech/Sockeye), the recently released ONT software for LR scRNA-seq analysis that also generates a cell barcode whitelist from nanopore long-reads.

### BLAZE identifies high confidence cell barcodes

Maximising sequencing depth per cell is key to accurately identifying and quantifying isoforms in single-cell data [3]. Therefore, we first compared the performance of BLAZE to Cell Ranger and Sockeye in the higher-depth PromethION data set. Cell Ranger, BLAZE and Sockeye identified 1022, 804 and 1518 cell barcodes respectively (**Table 2**). A comparison of barcodes showed 99.4% of barcodes identified by BLAZE were also found by Cell Ranger and Sockeye. However, a significant proportion of barcodes were unique to Cell Ranger and Sockeye (**Fig. 1C**). Analysis of cell barcode rank plots revealed BLAZE cell-associated barcodes had high read support in all methods (**Fig. 1D**). In contrast, unique Cell Ranger barcodes were often supported by few long reads, regardless of the different strategies of counting barcodes in BLAZE and Sockeye, suggesting that some barcodes identified by SR sequencing were not well represented in the LR dataset. Similarly, many unique Sockeye barcodes had little or no SR support, suggesting they are unlikely to be associated with cells. In addition, BLAZE counts for unique Sockeye barcodes were much lower (median 4.5 fold) than for barcodes found by both methods, suggesting many of the long reads supporting unique Sockeye barcodes were low quality and the barcodes are likely to be false positives.

**Table 2:**
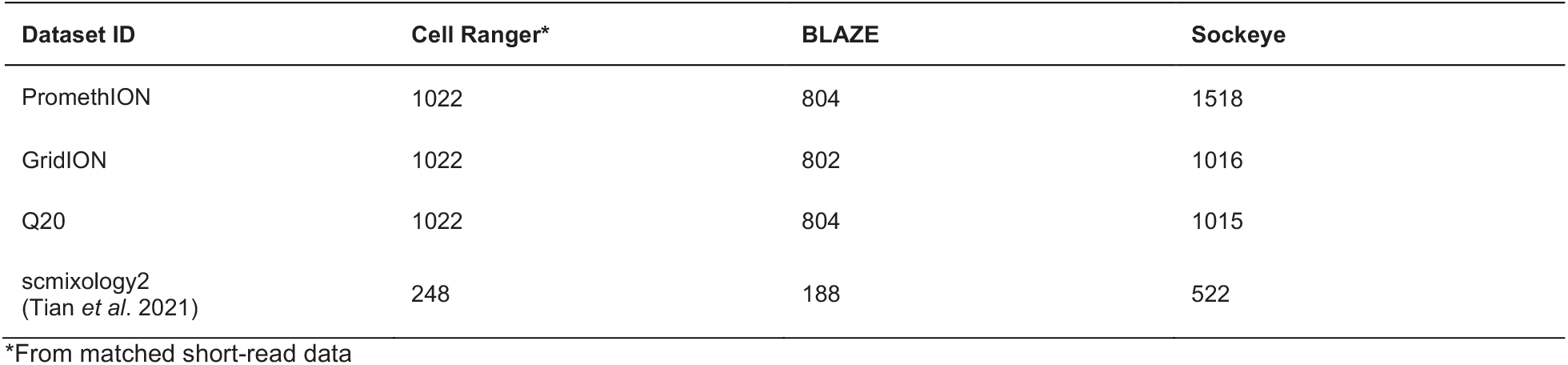
Number of barcodes detected.

| Dataset ID | Cell Ranger* | BLAZE | Sockeye |
| --- | --- | --- | --- |
| PromethION | 1022 | 804 | 1518 |
| GridION | 1022 | 802 | 1016 |
| Q20 | 1022 | 804 | 1015 |
| scmixology2<br>(Tian *et al.* 2021) | 248 | 188 | 522 |

\*From matched short-read data

The cell barcodes identified by Cell Ranger, BLAZE and Sockeye enabled the downstream analysis of a very similar proportion of reads, 70%, 69% and 69% respectively (useable reads, **Table 1**), demonstrating that the smaller number of barcodes found by BLAZE does not negatively affect the overall proportion of reads that can be assigned to a cell. Together these results show BLAZE provides the most accurate list of LR cell barcodes with little loss of sensitivity.

### Cell Ranger and Sockeye identify barcodes that are poorly supported by long-reads

We next asked if the barcode whitelists produced by Cell Ranger, BLAZE and Sockeye would yield similar results when clustering cells based on gene or isoform expression. We used the barcode whitelists and the ∼62 million long-reads from the PromethION as input into FLAMES [15] to produce gene and isoform counts and then generated UMAP plots in Seurat [20]. To facilitate comparison between the methods we made each UMAP plot separately and then coloured each cell according to its assigned cluster using the Cell Ranger whitelist. This revealed both Cell Ranger and Sockeye identify an additional cluster not found by BLAZE. This result was consistent for analyses using either isoform (**Fig. 2A**) or gene (**Additional File 1: Fig.S2A**) counts and was further confirmed by re-colouring the cells based on the BLAZE clusters (**Additional File 1: Fig.S2B**). This cluster contained poorly supported barcodes, as demonstrated by the low UMI counts and low numbers of genes and isoforms detected in each “cell” (**Fig. 2B and Additional File 1: Fig.S2C, D**). In the case of the Cell Ranger whitelist, these cells are likely those that exist in the matched SR data set, but are poorly represented amongst the long-reads, creating false positive long-read detections. The additional cluster found when using the Sockeye whitelist consists of a large number of cells not found with BLAZE or Cell Ranger and are also likely false-positives detections as they have low UMI and gene/isoform counts.

**Fig. 2:**
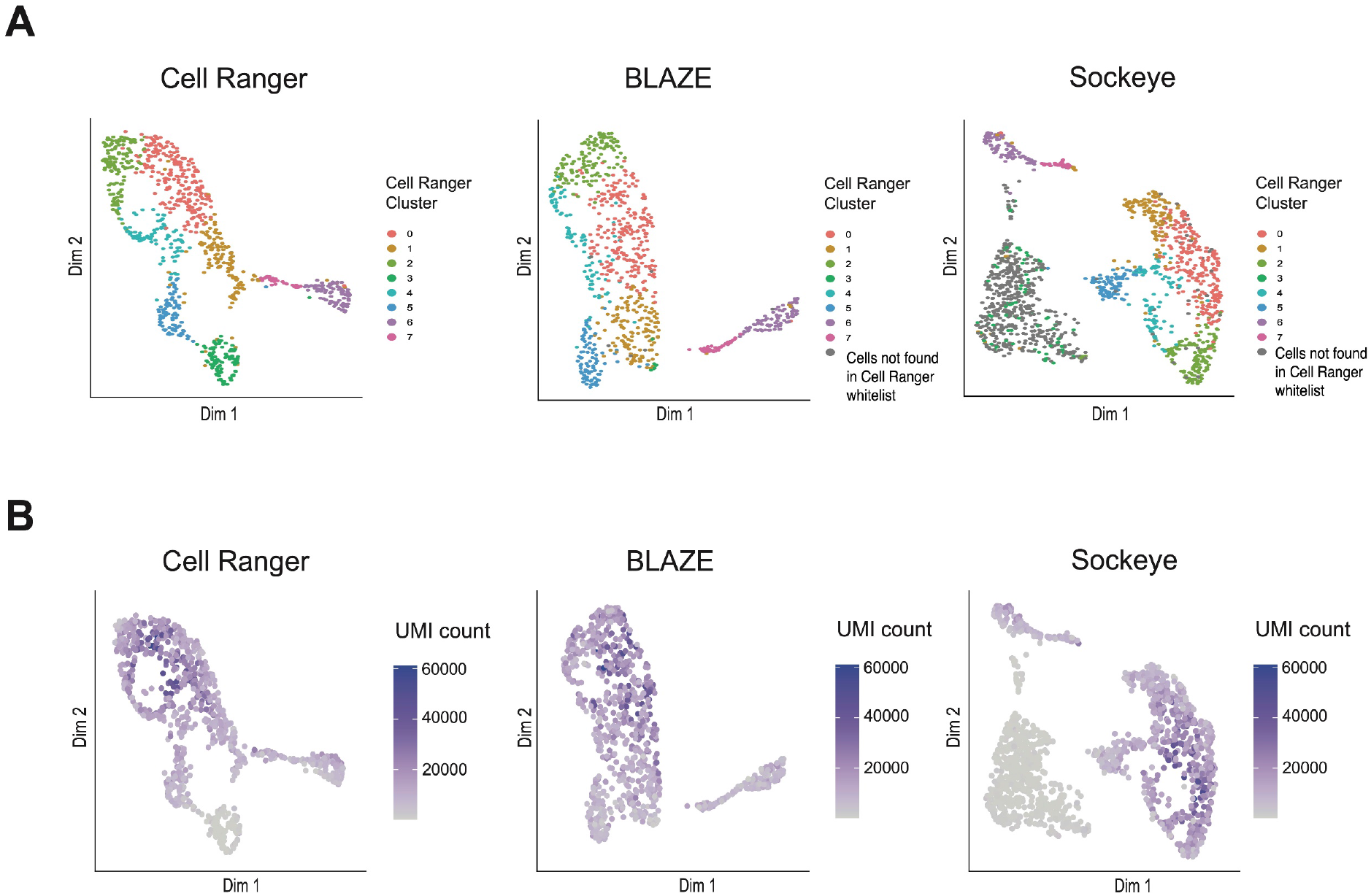
Comparison of cell clusters identified with BLAZE, Cell Ranger and Sockeye barcodes. Isoform expression UMAP plots from PromethION data. Isoform counts were generated with FLAMES using barcode whitelists from either Cell Ranger, BLAZE or Sockeye. **A:** Cells in all three plots are coloured based on clustering with the Cell Ranger whitelist. Cells not found in Cell Ranger whitelist are coloured in gray. **B:** Cells coloured based on UMI counts (sum of all unique UMIs across all transcripts) per cell.

The ∼1000 cells analysed here are in the early stages of cortical neuron differentiation, hence it was important to confirm BLAZE whitelist-based cell clustering was due to distinct biological profiles and not as a result of sequencing depth per cell or non-biologically relevant factors. Marker gene analysis confirmed biologically meaningful gene expression differences between clusters (**Additional file 2: Table S1 and Additional file 3: Table S2**). We identified significant differences in the expression of hundreds of genes including key transcription factors such as *ELAVL4* and *NHLH1* (**Fig. 3**), which are known to be upregulated during the differentiation of cortical neurons [21, 22]. Moreover, we find differential gene expression of well-defined neuron specific genes such as *NRN1* [23] and *PLPPR1* [24] (**Fig. 3**). Together these findings confirm that BLAZE cell clusters are transcriptionally distinct and that the BLAZE-FLAMES long-read pipeline is capturing the biological signal of neuronal cell differentiation.

**Fig. 3:**
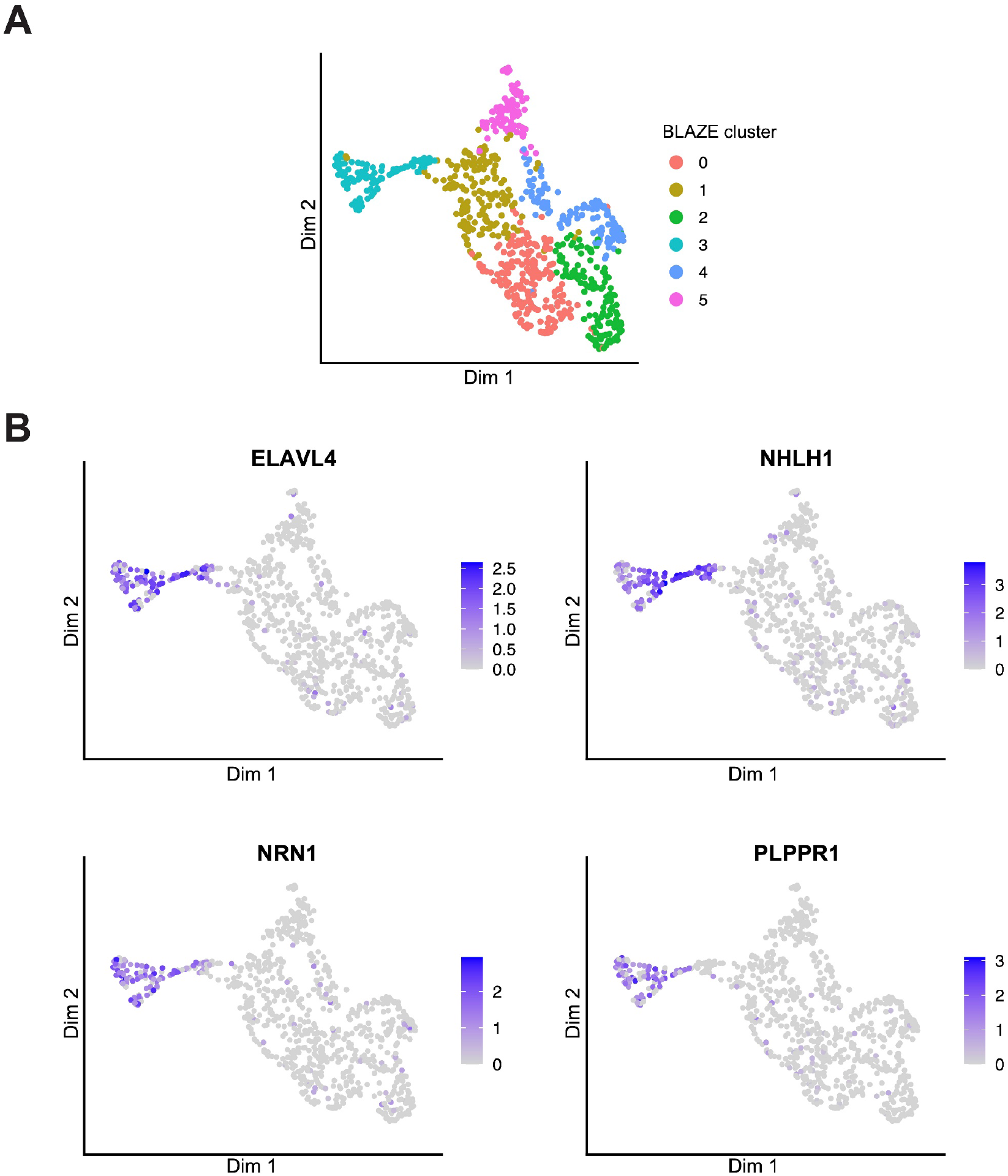
Gene expression UMAP coloured by cluster and expression of marker genes. **A:** UMAP showing clustering based on gene counts generated from FLAMES using BLAZE whitelist. **B:** UMAP coloured by expression of 4 marker genes known to be associated with differentiation and neuron development. Expression scale is coloured based on Seurat normalised counts. Colour scales are not comparable between plots.

While the use of FLAMES for isoform identification and quantification enables a fair comparison between whitelists, we wanted to ensure the false-positive detections from Sockeye were not a result of the FLAMES pipeline. To address this possibility, we implemented the complete Sockeye pipeline using default parameters and interrogated the UMAP plots generated by Sockeye. The Sockeye pipeline retained the additional cluster with low UMI counts (**Additional File 1: Fig.S3A**).We also note that Sockeye is currently limited to performing gene-based analyses and does not perform the isoform-based analyses enabled by LR scRNA-seq. Overall we find the BLAZE whitelist enabled the most accurate downstream expression and cell-type clustering of LR scRNA-seq data.

### Barcode detection with BLAZE is robust to changes in read depth or read accuracy

We investigated the impact of read depth and sequencing accuracy on the results of BLAZE. We sequenced the same single-cell cDNA sample on the lower-output Nanopore GridION, using both the LSK110 and higher accuracy Q20 chemistries. We find that although the LSK110 and Q20 GridION data produce significantly fewer total and pass reads compared to the PromethION (approximately 10% and 5% respectively) (**Table 1**), the number of barcodes found by BLAZE is virtually unchanged (**Table 2**). The Q20 GridION data is both lower depth and higher accuracy than the LSK110 data, leading to the possibility that higher read accuracy, (via an increased proportion of high confidence barcodes) could be maintaining barcode numbers. However, downsampling the LSK110 GridION data to match the Q20 read depth returned the same number of barcodes (802), demonstrating BLAZE performs consistently across data sets with variable read depths and different sequencing accuracies. In addition, we observed a similar proportion of usable reads between datasets (**Table 1**), implying that the improved Q20 accuracy had minimal effect on the number of reads that can be assigned to a cell.

We also assessed if Sockeye performed consistently across data sets of varying read depths. Sockeye identified 1016 and 1015 barcodes for LSK110 GridION and Q20 datasets respectively (**Table 2 and Additional File 1: Fig.S4**), which was a significant reduction on the 1518 barcodes from the PromethION data. UMAP results based on FLAMES quantification for the lower depth LSK110 and Q20 datasets revealed similar clustering between methods (**Fig. 4**). The number of barcodes detected by Sockeye (and subsequent downstream results) are therefore heavily dependent on per-cell read depth, leading to inconsistent results, with worse performance at higher read depths where isoform profiling is enhanced.

**Fig. 4:**
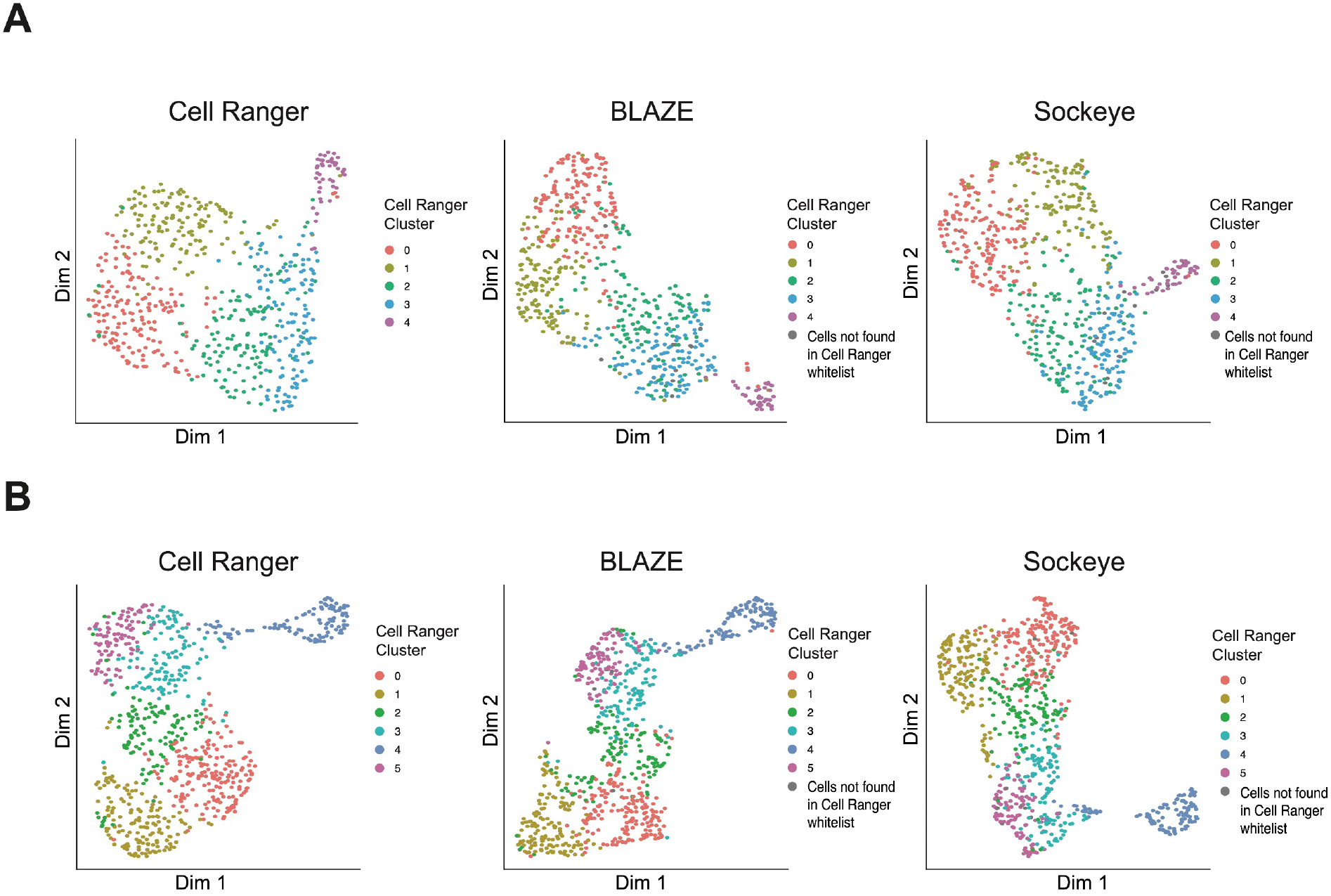
Isoform expression UMAP plot from Q20 and GridION data. **A:** Q20 **B:** GridION LSK110. Isoform counts were generated with FLAMES using barcode whitelists from either Cell Ranger, BLAZE or Sockeye. Cells are coloured as per Figure 2A.

We again tested if the full Sockeye pipeline would provide improved results over using the Sockeye barcodes in FLAMES. In contrast, we find that irrespective of the sequencing library used, quantification and UMAP generation using the Sockeye pipeline clusters cells in large part based on total UMI counts (**Additional File 1: Fig.S3**). A UMI associated clustering effect could potentially represent a real biological signal if it related to cells undergoing differentiation and changing their transcriptional activity. However, using the BLAZE-FLAMES-Seurat pipeline (instead of the complete Sockeye pipeline), we do not see such strong correlations between clusters and UMIs (**Additional File 1: Fig.S5**). These findings confirm the Sockeye pipeline is impacted by UMI associated confounders which bias UMAP results.

### BLAZE correctly identifies barcodes in long read single-cell data of known cell lines

To further validate the performance of BLAZE we compared Cell Ranger, BLAZE and Sockeye on an additional LR single-cell data set containing known and distinct cell lines. We utilised the scmixology2 data from Tian *et al*. (2021), which contains equal mixes of five cancer cell lines (∼40 cells per line) profiled with matched Illumina and Nanopore reads. Cell Ranger (from the matched short-reads), BLAZE and Sockeye identified 248, 188 and 522 cell barcodes respectively (**Table 2**). Similar to the cortical differentiation dataset we find all barcodes identified by BLAZE were also found by Cell Ranger and Sockeye (**Fig. 5A**). There were 59 barcodes identified by Cell Ranger and Sockeye but not by BLAZE and 275 barcodes unique to Sockeye **(Fig. 5A)**. These results suggested that Cell Ranger and Sockeye may be consistently identifying false positive long-read barcodes.

**Fig. 5:**
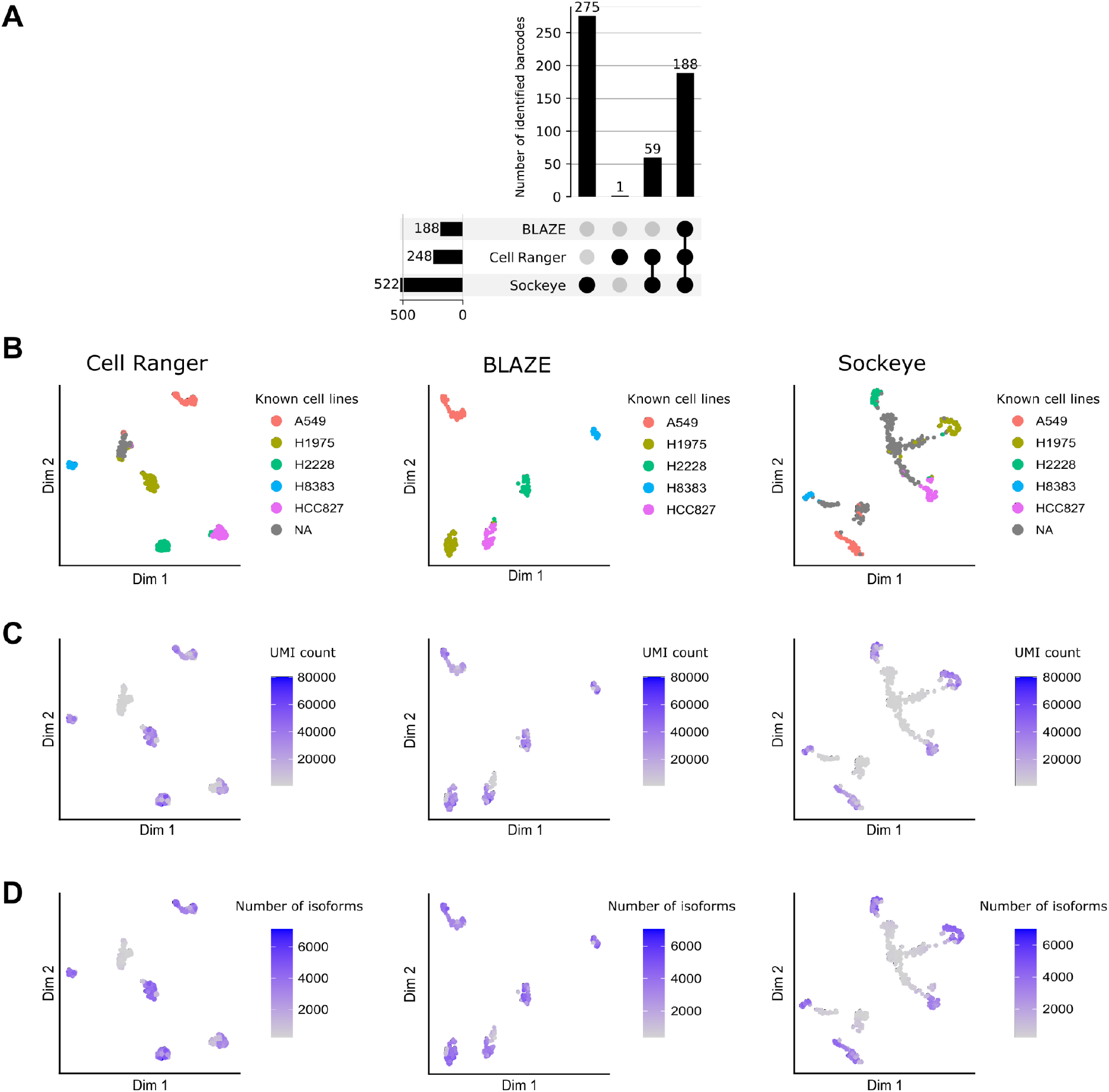
Barcode identification and clustering of Scmixology2 data. **A: Barcode upset plot comparing different whitelists.** Bar chart on left shows total number of barcodes found by each tool. Bar chart on top shows number of barcodes in the intersection of whitelists from specific combinations of methods. The dots and lines underneath show the combinations. **B-D: Isoform expression UMAP plots:** Isoform counts were generated with FLAMES using a barcode whitelist from either Cell Ranger (left), BLAZE (middle) or Sockeye (right). Cells are coloured based on: known cell types from Tian *et al*. 2021 **(B)**; Total UMIs per cell **(C)**; Number of isoforms detected in each cell **(D)**.

Implementation of the FLAMES pipeline for gene and isoform quantification supported the accurate identification of barcodes by BLAZE and confirmed the existence of false-positive detections by Cell Ranger and Sockeye (**Fig. 5B-D**). Scmixology2 contained five distinct cell lines and Tian *et al*. identified the barcodes belonging to each cell line in the LR data (see Tian *et al*. 2021 for details). We overlaid this information onto the UMAP plots generated from long-reads (**Fig. 5B**). UMAP plots generated from BLAZE barcodes detected the five expected cell lines. All cells found by BLAZE were present in the matched SR data (**Fig. 5A**), supporting the assertion that BLAZE accurately identifies cell barcodes while minimising false-positive detections. In contrast, Cell Ranger identified six distinct clusters. Five corresponded to the cancer cell lines in this sample **(Fig. 5B)**, while the sixth cluster, (denoted as N.A) largely comprised barcodes with no cell line match. These barcodes had very low cellular UMI counts and few unique isoforms (**Fig. 5C, D**) and likely represent cells present in the SR but not the LR data.

Clustering based on the Sockeye whitelist also identified additional cell type clusters, with the majority (52%) of cells in clusters not matching one of the known cell lines. These “cells” all have low UMI counts and fewer detected isoforms (**Fig. 5C, D**), highlighting that these barcodes likely represent false positives and are not real cells. To ensure these findings were not a consequence of the FLAMES pipeline we also ran the entire Sockeye workflow. The Sockeye generated UMAP displayed similar results (**Fig. S6**), further supporting incorrect barcode identification by Sockeye. The identification of false positive barcodes and cell clusters when using the Cell Ranger and Sockeye whitelist again demonstrate that BLAZE produces a more accurate representation of barcodes present in LR datasets.

### Overall comparison between BLAZE and Sockeye

BLAZE is more conservative than Sockeye in calling barcodes and therefore minimises false-positive detections. However, both BLAZE and Sockeye use barcodes with counts above a threshold to generate the whitelist and users have the flexibility to choose the count threshold to trade off high precision (i.e. fewer false barcodes) for high recall (i.e. more true barcodes). Using all four datasets above (**Tables 1 and 2**) and defining the cell barcodes identified by Cell Ranger as the ground truth, we calculated precision-recall curves across different count thresholds in BLAZE and Sockeye. Results demonstrated that BLAZE consistently outperforms Sockeye (**Fig. 6**) and outputs a better whitelist regardless of whether users prefer high precision or recall.

**Figure 6:**
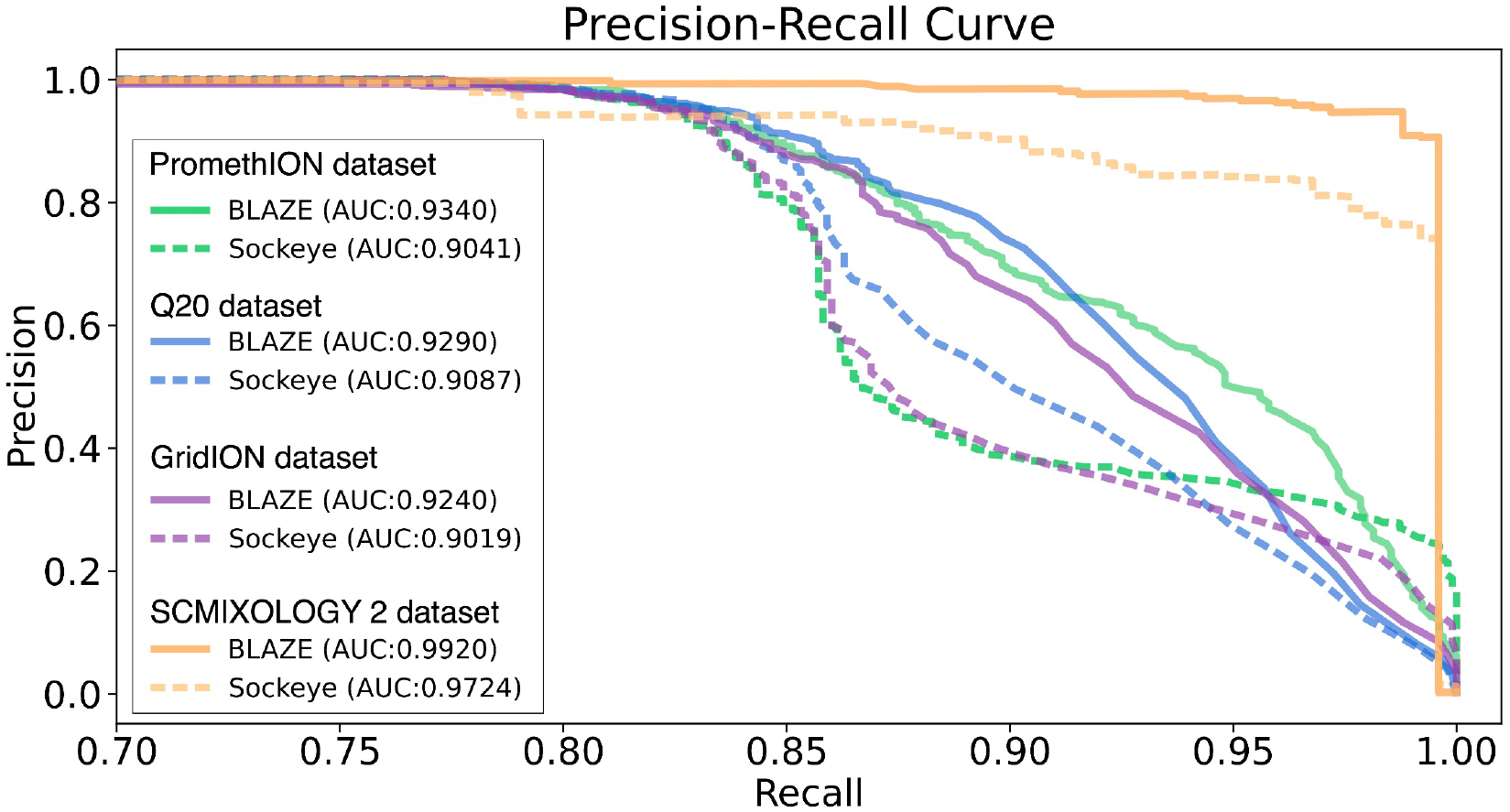
Precision-recall curves across different datasets for BLAZE and Sockeye. Precision and recall were calculated across different count thresholds by using the barcodes identified from short reads (i.e. whitelist from Cell Ranger) as the ground truth. The numbers in the legend show area under the curve (AUC) values.

BLAZE is easy to install and run (see **Additional File 4: Tables S3** for the runtime of BLAZE). However, a fair runtime comparison between BLAZE and Sockeye is difficult because Sockeye is not designed to solely generate a barcode whitelist but instead runs the whole pipeline for single-cell gene expression and therefore requires a longer runtime. In addition, Sockeye cannot be utilised as a stand-alone tool to perform single-cell isoform analysis, (for which long-reads are significantly more useful than short-reads) as it only performs gene-level quantification. In this sense, running BLAZE is quicker and the integration is easier as BLAZE outputs a whitelist using the Cell Ranger format that can be input into tools such as FLAMES without modification.

## Discussion

Single-cell RNA sequencing (scRNA-seq) has revolutionised the study of transcriptomes, yet is limited by the use of SR sequencing methods. With recent advancements in LR scRNA-seq methodologies [5, 25] and analysis tools [26], the potential to study the complete array of RNA isoforms and quantify isoform expression at single-cell resolution is becoming possible. The use of “noisy” long-reads however, presents its own unique set of challenges, primarily the difficulty in identifying the cell barcodes needed to assign each transcript to its cell of origin. Consequently, the use of matched SR data has been fundamental to the successful implementation of high depth, high throughput nanopore LR scRNA-seq. In spite of the higher error rate of nanopore reads, we show that BLAZE aids in eliminating the need for matched SR sequencing. This not only simplifies the procedure but also reduces overall library construction and sequencing costs and therefore increases the accessibility of LR scRNA-seq.

We found BLAZE to be robust in its ability to accurately identify 10x cell barcodes from long-reads. BLAZE can be applied to different types of single-cell samples and performs equally well on both higher accuracy Q20 data, as well as lower accuracy reads generated from ONT’s LSK110 and LSK109 protocols. We find that ONT’s recently published software for long-read only barcode identification, Sockeye, appears to be affected by read-depth associated confounders and identifies false-positive cell barcodes. An alternate possibility is that Sockeye is more effective than BLAZE at identifying cell barcodes and therefore finds larger numbers of cells. However, this seems unlikely given Sockeye finds much larger numbers of long-read barcodes than matched short-read sequencing; unique Sockeye barcodes don’t match the known cell types present in the scmixology2 data; and the unique “cells” have very low numbers of UMIs, genes and unique isoforms. In order to accurately identify and quantify isoforms from scRNA-seq it is important to sequence cells deeply [3]. BLAZE showed the greatest advantage over other methods in the higher depth PromethION datasets and therefore performs well in the context most relevant to LR scRNA-seq.

We designed BLAZE to be simple to install and use and seamlessly integrate into existing isoform identification and quantification pipelines such as FLAMES, meaning no modifications to existing protocols or pipelines are needed. This provides a further advantage over Sockeye, which currently only performs gene level quantification. Most importantly and perhaps unexpectedly, we find that BLAZE outperforms barcode whitelists generated from matched SR data using Cell Ranger. More than 99% of barcodes identified with BLAZE were present in the SR whitelist confirming that false-positive detections with BLAZE are rare. Conversely, >20% of barcodes identified by Cell Ranger were not found by BLAZE. These barcodes were supported by few long-reads and expressed comparatively fewer genes and isoforms. We hypothesise that despite sequencing matched samples some cell barcodes found in SR data are poorly represented amongst the long-reads. Supporting this, the Cell Ranger knee plot showed the barcodes not found by BLAZE had low UMI counts in the SR data. Such barcodes are the most likely not to be found in matched LR sequencing due to chance and differences in read depths. Consequently, the use of long-read only barcode identification methods should produce whitelists that more faithfully represent cells profiled with long-read sequencing.

The accurate identification of single-cell barcodes is crucial to downstream gene and isoform quantification. Nearly all single-cell workflows cluster cells based on expression using dimensional reduction techniques such as t-SNE [27] and UMAP [28, 29]. These methods enable further integration of cell type specific markers and can be used to identify differentially expressed genes and isoforms between cell clusters. False-positive cells often cluster together giving a misleading impression of additional cell clusters, which could confound differential expression analyses and biological interpretation of the results. Furthermore, usable reads can be assigned to false-positive barcodes, reducing the read depth of real cells and decreasing experimental power for isoform identification and quantification. Filtering out cells that have low UMI counts could reduce false-positive cells, however deciding on an appropriate UMI filtering threshold can be difficult and would depend on sequencing read depth and the transcriptional activity of the cells. It can be challenging to distinguish between cells that produce small amounts of RNA (and subsequently have few UMIs) and false-positive cells. Tools designed to generate single-cell barcode whitelists should therefore prioritise high precision as false-positive barcodes can confound downstream workflows.

A limitation with the current study is the use of Cell Ranger as the ground truth to determine the precision-recall of BLAZE and Sockeye, since our results suggest some barcodes identified by Cell Ranger do not represent genuine cells in the LR data. This acts to decrease the recall of BLAZE, while inflating the precision of Sockeye. Even so, we find BLAZE precision-recall systematically outperforms Sockeye and we conclude the outperformance would be even greater with a perfect ground truth dataset.

Currently BLAZE is limited to identifying 10x single-cell barcodes from nanopore reads. Although other LR single-cell methodologies such as scCOLOR-seq [13] and R2C2 [30] have been used to profile single cells with long-reads, the 10x chromium platform is the most widely available and popular platform. We therefore designed the initial version of BLAZE to facilitate 10x barcode identification. Recent developments in throughput and accuracy for PacBio HiFi sequencing are increasing the applicability of PacBio for LR scRNA-seq, while LR nanopore protocols for other scRNA-seq modalities such as Split-seq are also now available [14, 31-34]. Although BLAZE is currently limited to identification of 10x barcodes from nanopore reads, we see potential to expand BLAZE to process both PacBio HiFi reads and reads from other scRNA-seq methods in the future.

## Conclusion

We show that BLAZE is a highly accurate single-cell barcode identification tool for Nanopore long-reads. We demonstrate that BLAZE works well across different data sets, read depths and read accuracies and can seamlessly integrate into existing tools for downstream gene and isoform identification and quantification. Crucially, BLAZE eliminates the requirement for additional matched SR data and therefore simplifies LR scRNA-seq protocols while significantly reducing cost. BLAZE has been designed to be widely accessible and easy to use and is available at https://github.com/shimlab/BLAZE.

## MATERIALS AND METHODS

### Cell lines and Stem Cell Differentiation

RM3.5 human induced pluripotent stem cells (hiPSC) [35] were cultured under xenogeneic conditions in accordance with the protocol described in Niclis *et*.*al* [36]. PSCs were differentiated into cortical neuron lineage using the protocol described by Gantner *et*.*al*. [37].

### Preparation of single-cell suspension

At day 26 post neural induction RM3.5 cells undergoing cortical differentiation were harvested for analysis. Cells were washed twice in 300 mL of DPBS -/-and exposed to Accutase (Innovative Cell Technologies, Inc. San Diego, CA, http://www.accutase.com) for 12 min at 37°C. Following incubation, cells were moved to a 15 mL falcon tube and were gently triturated to help generate a single-cell suspension. DPBS was added at 1:1 ratio to inactivate the Accutase and the sample gently centrifuged at 1500 rpm for 3 min at 4 °C and supernatant removed. Cells were resuspended in 2 mL DBPS and Rock inhibitor Y-27632 (diluted 1:1000) (Tocris Bioscience) to prevent cell death. The cell suspension was passed through a Flowmi™ strainer (Flowmi; Cat. No. 64709-60) to remove remaining cell debris. Finally, cells were counted using a hemocytometer and viability assessed with trypan blue stain (ThermoFisher scientific Cat. No. 15250061) prior to final resuspension in DPBS with 0.04% BSA and Rock inhibitor.

### FLT-seq 10x single-cell processing and cDNA amplification

FLT-seq was performed in accordance with the published protocol ([15], https://www.protocols.io/view/massively-parallel-long-read-sequencing-of-single-81wgbpp1nvpk/v1).

Briefly, the cell suspension was prepared for target recovery of 5000 cells, with 20% for matched short and long-read sequencing. Single-cell processing and cDNA amplification was performed in accordance with the 10x Genomics Chromium Single-cell 3’ gene expression protocol (v3.1), except that to generate full-length cDNA reverse transcription the extension time was extended to 2 hours. GEMs were split 80%:20%, with the cDNA from the 20% (∼1000 cells) processed to create matched short and long-read libraries. We used FLT-seq as this protocol generates a high proportion of full-length (3’ adaptor to 5’ TSO) reads and an almost negligible proportion of TSO artifacts (TSO-TSO reads without a valid cell barcode).

### Short-read Illumina sequencing

The Illumina short-read library was sequenced on the Novaseq6000 to a depth of 100 M reads. Base calling and quality scoring were determined using Real-Time Analysis on board software RTA3, while the FASTQ file generation and de-multiplexing utilised bclConver v3.9.3.

### Nanopore single-cell library preparation and sequencing

Full length cDNA generated from the FLT-seq protocol was prepared using the SQK-LSK110 Ligation Sequencing Kit (ONT) with the following modifications: incubation times for end-preparation and A-tailing were lengthened by 15 min and all AMPureXP cleaning steps were performed at x1.8. Libraries were sequenced on both the GridION (FLO-MIN106 flow cell) and PromethION (FLO-PRO002 flow cell) loading ∼45 fmol with an additional flow cell top up with any remaining library at 24 hrs. Fast5 files were generated using MinKnow v21.02.5 on the GridION and v22.03.4 on the PromethION and basecalled with guppy v6.0.7 with the super high accuracy configuration file.

We prepared an additional long-read library with the SQK-Q20EA Genomic DNA by ligation Q20+ early access kit (ONT) with the same modifications stated above. We sequenced the Q20 library on the GridION (FLO-MIN112 flow cell), loading 10 fmol with an additional 10 fmol top up at 24 hrs. Fast5 files were generated using MinKnow v21.05.25 and basecalled with guppy v6.0.7 with the dna_r10.4_e8.1_sup.cfg configuration file.

Median sequencing accuracy was calculated by first mapping pass FASTQ files to the transcriptome with Minimap2 [38] using the command minimap2 -ax map-ont $REF $FASTQ > trans_mapping.sam. Median accuracy was calculated using a custom R script found at https://github.com/josiegleeson/BamSlam [39]. In short, the cigar strings from primary alignments were extracted and the total number of mismatches and insertions and deletions per alignment were calculated.

### Identification putative barcode sequence in each read

BLAZE identifies the likely position of the cell barcode (referred to as “putative barcode”) by first identifying the position of the adaptor. Similar to [9], in each nanopore read, BLAZE searches for the last 10 nt sequence of the adaptor (i.e. “CTTCCGATCT”) in the first 200 nt of the read. Specifically, BLAZE aligns the “CTTCCGATCT” to the first 200 nt of the read using Biopython [40] and allows up to 2 mismatches, insertions or deletions. This procedure ensures a high sensitivity in identifying the adaptor location but will potentially find multiple locations. Thus, BLAZE also requires a downstream polyT sequence for accurate identification of the adaptor location. Specifically, BLAZE conducts a lenient search that looks for 4 consecutive ‘T’s 20∼50 nt downstream of the adaptor, as the polyT tail in nanopore reads is often truncated due to limitations in basecalling of homopolymers [12]. The corresponding adaptor is considered to be valid only if the polyT is found. BLAZE then repeats the same procedure for the reverse complement sequence. Reads with exactly 1 valid adaptor were kept for the downstream steps. The 16 nt sequence immediately downstream of the adaptor is defined as the “putative barcode”.

### Selection of high-quality putative barcodes

To accurately identify the sequences of barcodes, BLAZE selects high-quality putative barcodes that are less likely to contain basecalling errors. Basecalling outputs provide a (Phred) quality score for each base, which indicates the probability of the base being correctly basecalled. Incorrectly basecalled bases generally have a low quality score, so putative barcodes with error(s) are more likely to have at least one base with a low quality score. Therefore, for each putative barcode, BLAZE calculates the minimum of quality scores across the 16 bases in the putative barcode, denoted as “minQ”, and selects putative barcodes with minQ ≥15 as high-quality putative barcodes. See **Figure S1** for our choice of 15 as a threshold.

### Identification of cell-associated barcodes from high-quality putative barcodes

BLAZE lists unique high-quality putative barcodes, counts their occurrences, and ranks them based on those counts. Next, similar to Zheng et al 2017, BLAZE selects those barcodes whose counts are larger than a stringent count threshold *T* as cell-associated barcodes (i.e., barcodes likely associated with cells), and outputs them in a whitelist. The threshold *T* has been chosen as follows. For a given expected number of recovered cells, denoted by *N*, we obtain c, the count of a unique high-quality barcode whose rank is 0.95×*N*. Then, we use 0.05×*c* as the threshold *T*. In practice, the targeted number of cells can be a plausible number for *N*. We use *N =* 500 in the analysis in this manuscript. The number of barcodes in a final whitelist is robust to the choice of *N* (**Figure S7**) as long as N is set within a reasonable range that is not too divergent from the true number (e.g., the number of barcodes change from 186 to 193 when N is increased from 50 to 1500 in the analysis of the scmixology2 dataset with ∼200 cells).

### Barcode whitelist generation and gene and isoform qualification with FLAMES

We produced barcode whitelists using three software packages. Cell Ranger v6.0.2, Sockeye v0.2.1 (ONT) (https://github.com/nanoporetech/Sockeye) and BLAZE v1.0.0 (https://github.com/shimlab/BLAZE). First, we processed fastq files generated from the matched Illumina sequencing using the Cell Ranger pipeline to generate the barcode whitelist. Next, we ran the Sockeye pipeline and BLAZE on each long-read data set using default parameters to generate barcode whitelists from long-reads only. We performed gene and isoform level qualification using FLAMES [15] (https://github.com/OliverVoogd/FLAMES) using an edit distance of 2, hg38 reference genome and GENCODE v31 comprehensive transcriptome. We used isoform count matrices generated by FLAMES to produce gene level counts using a custom python script (available at https://github.com/youyupei/bc_whitelist_analysis/).

### UMAP generation and single-cell data processing

Gene and isoform count matrices were analysed with the R package Seurat v4.1.1 [20]. We applied a minimum filtering threshold of 200 features (genes or isoforms) to remove cells with very low UMI counts in accordance with Seurat pipeline recommendations. Clustering was performed on all data sets with a resolution value of 0.7. Marker genes/isoforms that distinguish clusters were found using Seurat::FindMarkers using default parameters, full workflow available at https://github.com/youyupei/bc_whitelist_analysis/blob/main/script/SC_Marker_gene.Rmd. Seurat analysis scripts and output files can be found at https://github.com/youyupei/bc_whitelist_analysis.

### Scmixology 2 data set

Fast5 files from the scmixology 2 data set published in Tian *et al*. (2021) were rebasecalled with guppy v5.1.13 to generate fastq files. We generated long-read barcode whitelists using BLAZE and Sockeye as stated above. The Cell Ranger generated whitelist was obtained from matched Illumina short-read sequencing published in Tian *et al*. (2021). These three whitelists were inputs into FLAMES for gene and isoform quantification and downstream processing with Seurat is as stated above.

### Ethics approval and consent to participate

All research activities involving iPSC lines were performed under institutional ethics approval from The University of Melbourne Ethics ID 1239208.

## Consent for publication

Not applicable.

## Availability of data and materials

Fast5 and fastq files are available from ENA under accession PRJEB54718. The processed data and scripts used in this study are available at https://github.com/youyupei/bc_whitelist_analysis/. BLAZE is implemented in Python and available on github at https://github.com/shimlab/BLAZE under GNU General Public License v3.0.

## Authors’ contributions

MBC conceived the study. YY wrote BLAZE. CH performed cell culture. CP oversaw cell culture. YP performed single-cell experiments and sequencing with assistance from RDP. YY and YP performed bioinformatic analyses. MBC and HS oversaw research. YY, YP, HS and MBC wrote the paper with input from all authors.

## Acknowledgements

The RM3.5 cell line was supplied by Prof. Ed Stanley, Murdoch Children’s Research Institute and Monash University. This research was undertaken using the LIEF HPC-GPGPU Facility hosted at the University of Melbourne. This Facility was established with the assistance of LIEF Grant LE170100200. **Figure 1B** was created under license with BioRender.com. The authors thank Chiara Pavan for expert technical assistance and the lab of A/Prof Matt Ritchie at WEHI for assistance running FLAMES, sharing sequencing data and helpful feedback on the study.

## Supplementary Information

**Additional file 1:** Fig. S1: Distribution of minQ. Fig. S2. UMAP plots from PromethION data. Fig. S3. Gene expression UMAP plot from Sockeye pipeline for PromethION, GridION and Q20 data. Fig. S4. Barcode upset plot comparing different whitelists. Fig. S5. Gene expression UMAP plot (using BLAZE whitelist) from PromethION, GridION and Q20 data. Fig. S6. Gene expression UMAP plot from Sockeye pipeline for scmixology 2 data. Fig. S7. Effect of specifying different numbers of expected cells in BLAZE.

**Additional file 2:** Table S1. Genes differentially expressed between BLAZE cell clusters

**Additional file 3:** Table S2. Genes differentially expressed in the differentiating cell cluster (BLAZE analysis)

**Additional file 4:** Table S3. BLAZE runtime (with 32 cpus)

