## Additional file 1 for "Identification of cell barcodes from long-read single-cell RNA-seq with BLAZE"

### Fig. S1

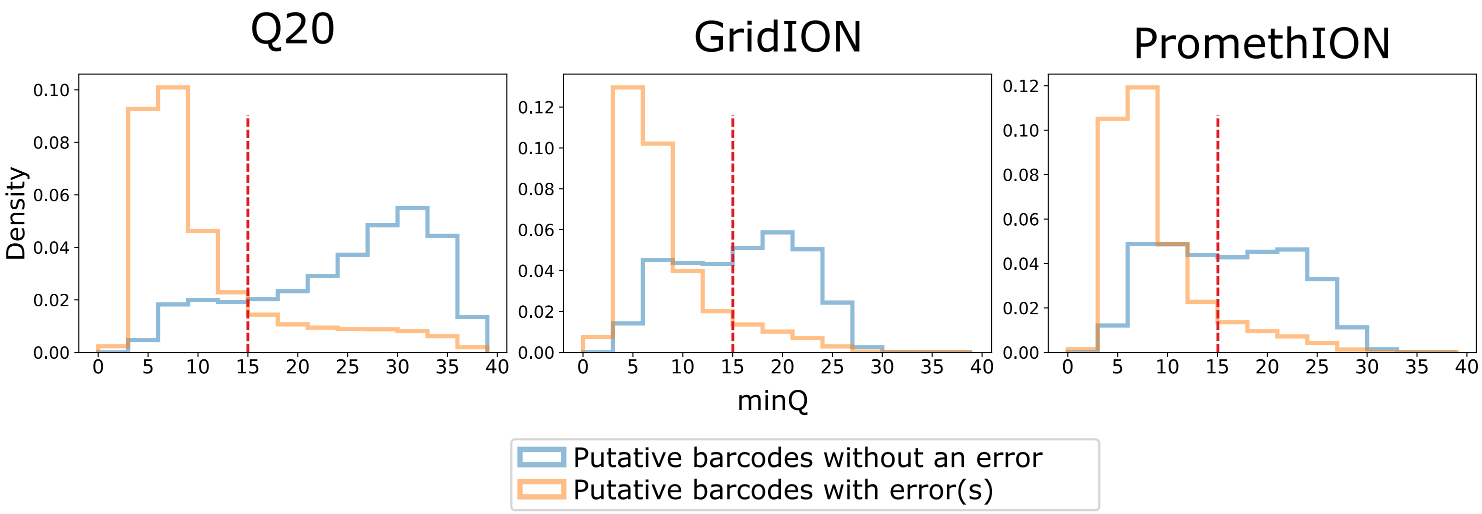


**Fig. S1: Distribution of minQ**: The title for each panel represents a different dataset in Table 1. Using the barcodes identified from short reads (by Cell Ranger) as a ground truth, all putative barcodes could be categorised as either: 1. Without error (exact match to ground truth barcode); 2. With error(s). While it's possible a putative barcode contains an error but exactly matches a ground truth barcode by chance, this should be very rare and not significantly impact the distribution. The red dotted lines indicates minQ=15, which is the threshold used in BLAZE to identify high-quality putative barcodes.

### Fig. S2

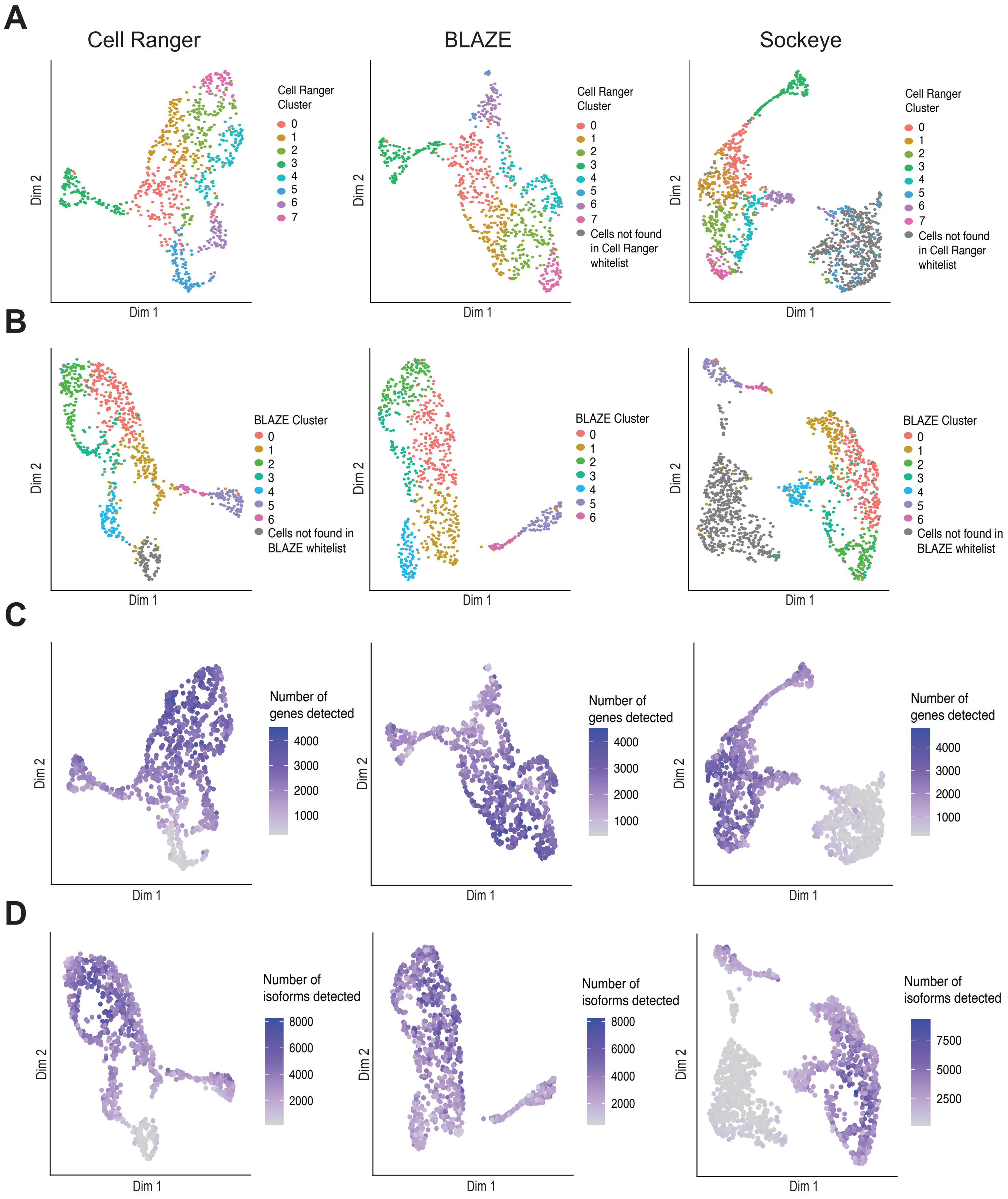


**Fig. S2. UMAP plots from PromethION data.** Counts were generated with FLAMES using barcode whitelists from either Cell Ranger, BLAZE or Sockeye. **A**. Gene expression UMAP coloured by Cell Ranger clusters: Cells in all three plots are coloured based on clustering with the Cell Ranger whitelist. Cells not found in Cell Ranger whitelist are coloured in gray. **B**. Isoform expression UMAP coloured by BLAZE clusters: Cells in all three plots are coloured based on clustering with the BLAZE whitelist. Cells not found in BLAZE whitelist are coloured in gray. **C**. Gene expression UMAP coloured by number of genes detected per cell. **D**. Isoform expression UMAP coloured by number of unique isoforms detected per cell: Unique isoforms are defined as structurally distinct RNA isoforms with a least one UMI count.

### Fig. S3

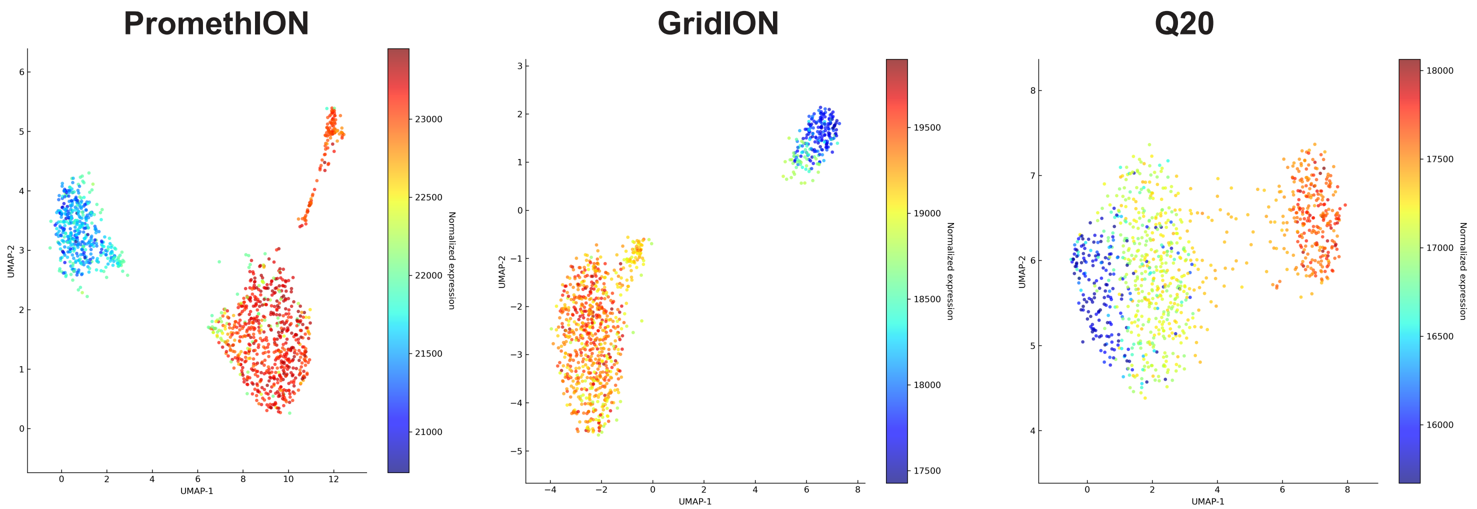


**Fig. S3. Gene expression UMAP plot from Sockeye pipeline (PromethION, GridION and Q20 data).** Cells are coloured based on normalised gene expression calculated within the Sockeye pipeline.

### Fig. S4

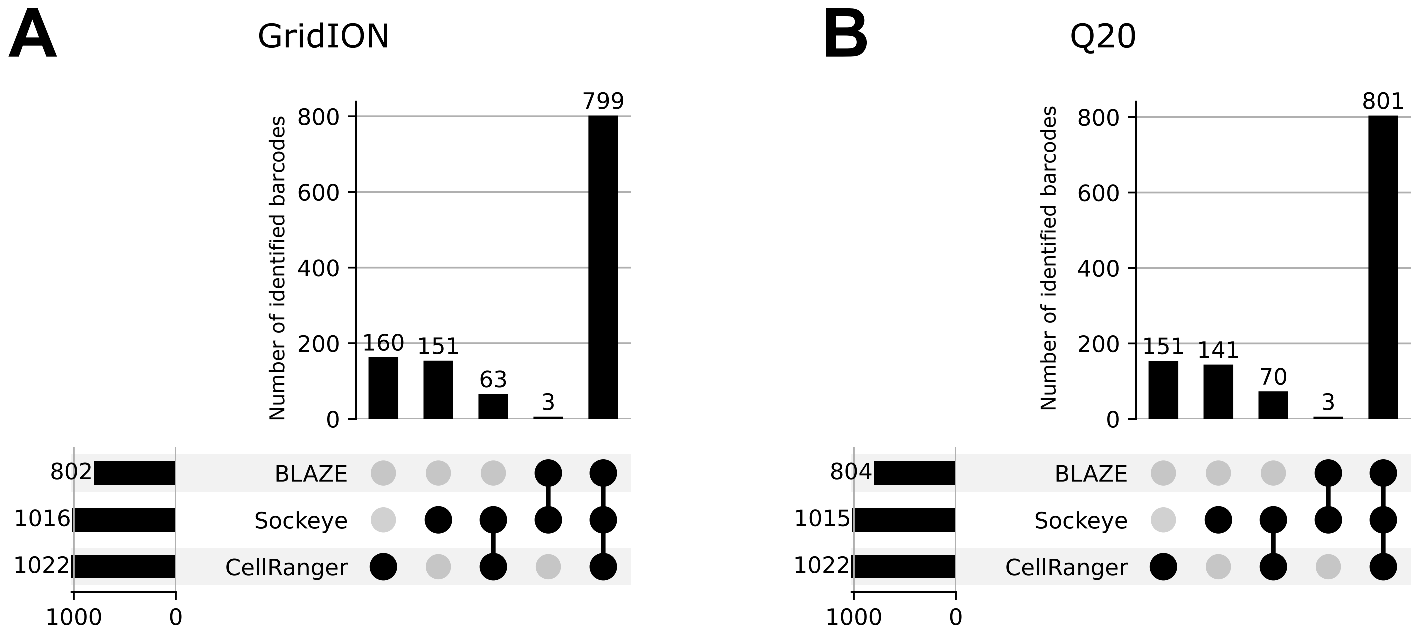


**Fig. S4. Barcode upset plot comparing different whitelists** in **A**. LSK110 GridION data and **B**. Q20 GridION data. The bar chart on the left shows the total number of barcodes found by each tool. The bar chart on the top shows the number of barcodes in the intersection of whitelists from specific combinations of methods. The dots and lines underneath show the combinations.

### Fig. S5

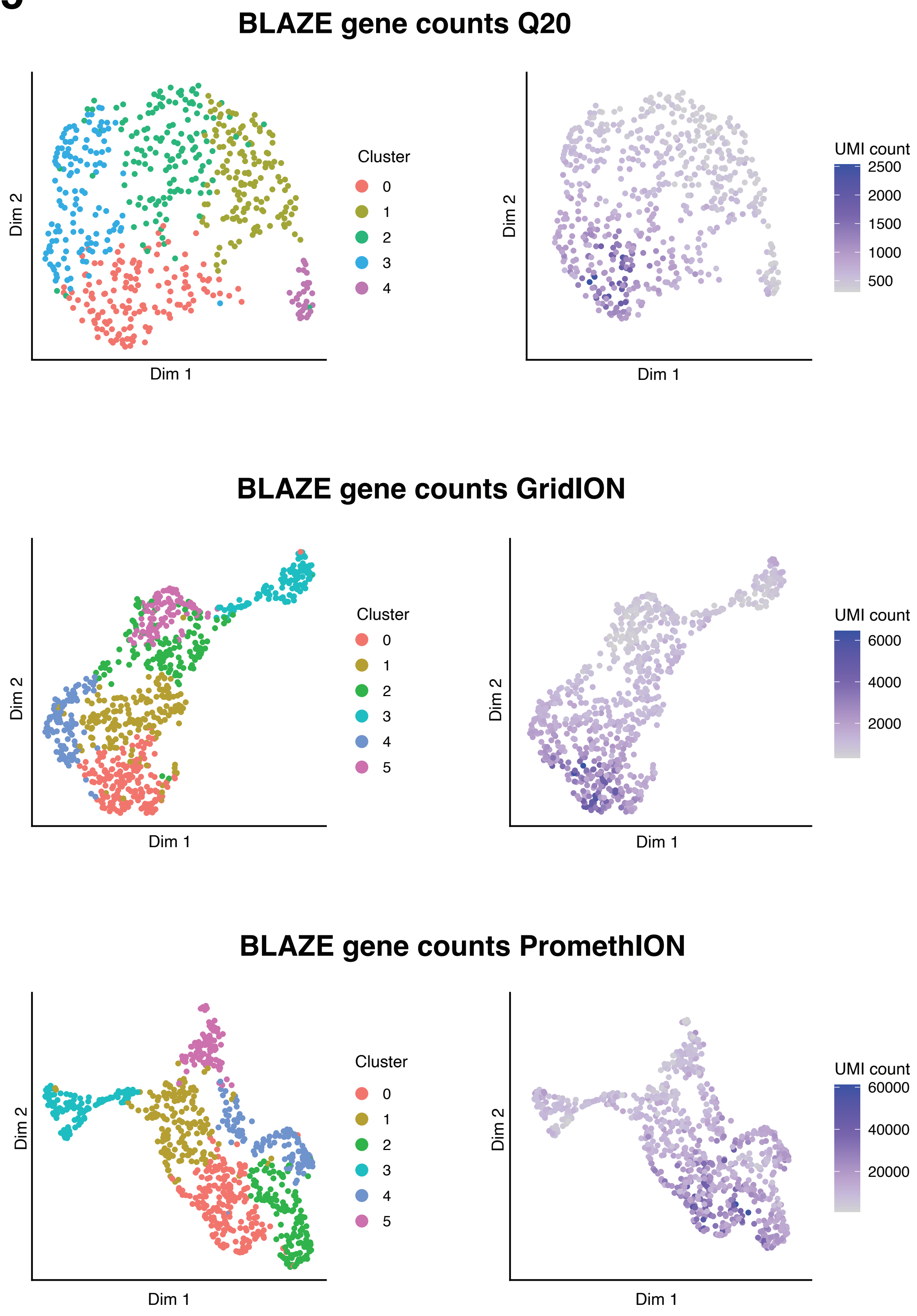


**Fig. S5. Gene expression UMAP plot (using BLAZE whitelist) from PromethION, GridION and Q20 data.** Gene expression is generated by FLAMES using BLAZE whitelist as input. Cells are coloured based on cluster (left) and UMI count (right).

### Fig. S6

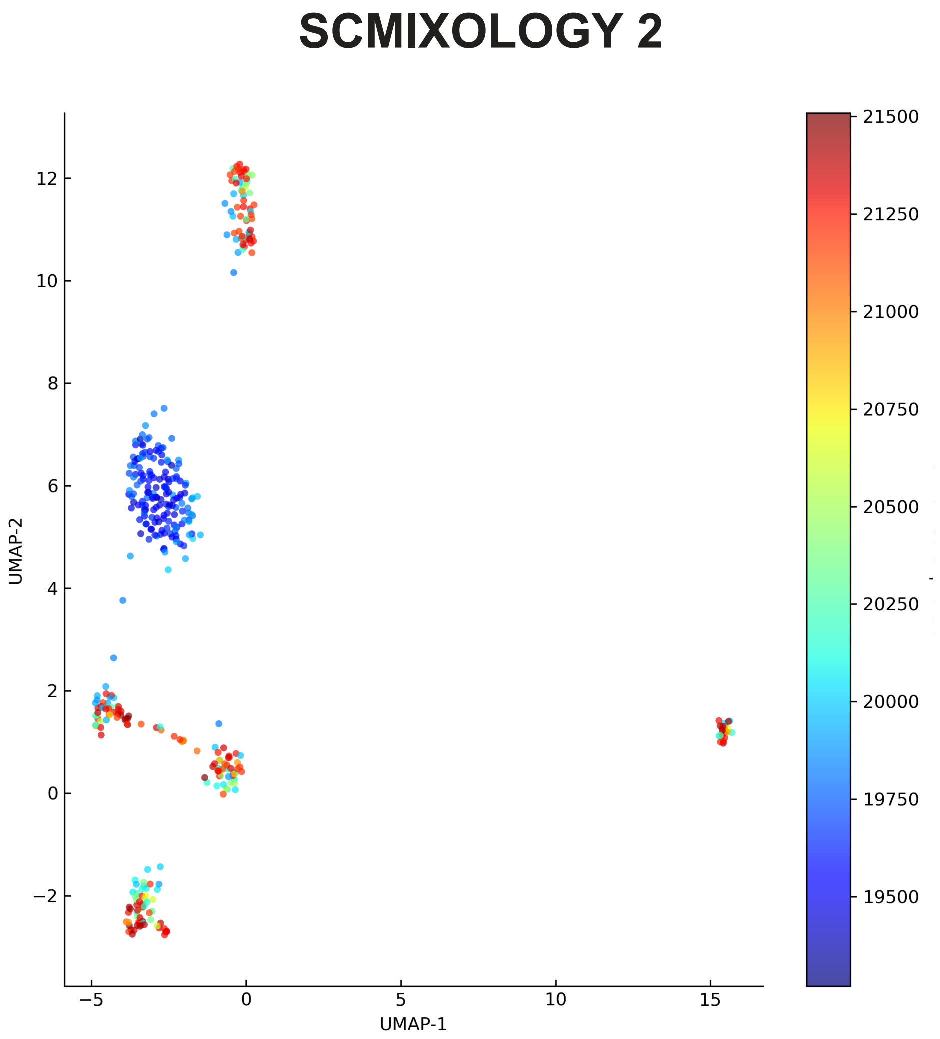


**Fig. S6. Gene expression UMAP plot from Sockeye pipeline (scmixology 2 data).** Cells are coloured based on normalised gene expression calculated within the Sockeye pipeline.

### Fig. S7

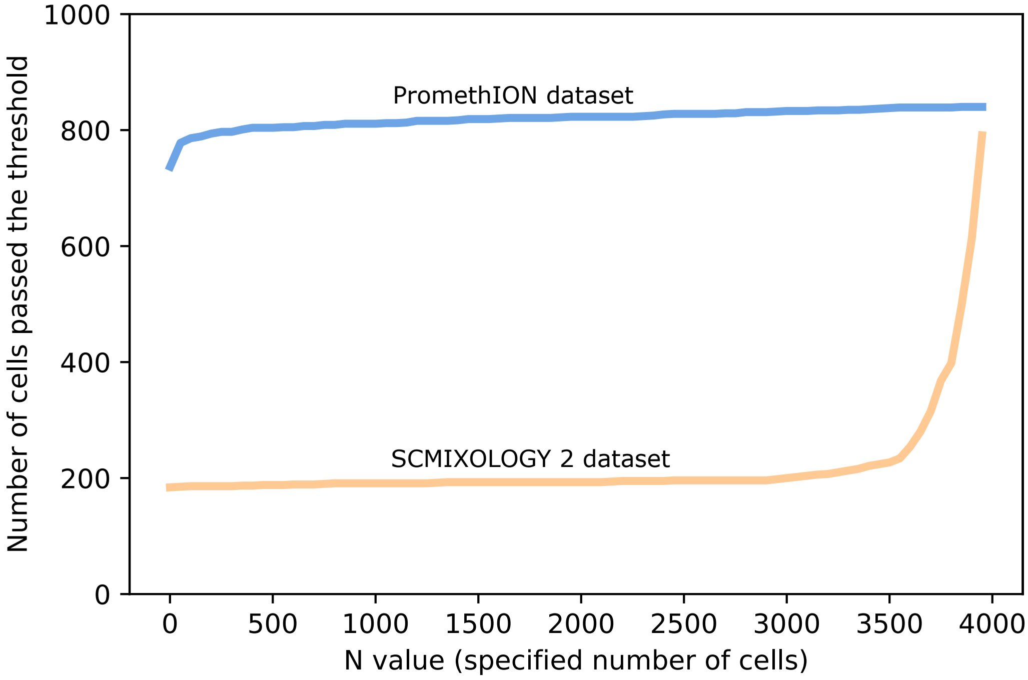


**Fig. S7. Effect of specifying different numbers of expected cells in BLAZE.** The count threshold in BLAZE is determined partly based on a specified number of cells (N). Figure shows how the number of cells passing this threshold (i.e. the identified number of cells) changes with different values of N in the PromethION and scmixology 2 datasets.
